# A Hidden Markov Model for Detecting Confinement in Single Particle Tracking Trajectories

**DOI:** 10.1101/275107

**Authors:** PJ Slator, NJ Burroughs

## Abstract

State-of-the-art single particle tracking (SPT) techniques can generate long trajectories with high temporal and spatial resolution. This offers the possibility of mechanistically interpreting particle movements and behaviour in membranes. To this end, a number of statistical techniques have been developed that partition SPT trajectories into states with distinct diffusion signatures, allowing a statistical analysis of diffusion state dynamics and switching behaviour. Here we develop a confinement model, within a hidden Markov framework, that switches between phases of free diffusion, and confinement in a harmonic potential well. By using a Markov chain Monte Carlo (MCMC) algorithm to fit this model, automated partitioning of individual SPT trajectories into these two phases is achieved, which allows us to analyse confinement events. We demonstrate the utility of this algorithm on a previously published dataset, where gold nanoparticle (AuNP) tagged GM1 lipids were tracked in model membranes. We performed a comprehensive analysis of confinement events, demonstrating that there is heterogeneity in the lifetime, shape, and size of events, with confinement size and shape being highly conserved within trajectories. Our observations suggest that heterogeneity in confinement events is caused by both individual nanoparticle characteristics and the binding site environment. The individual nanoparticle heterogeneity ultimately limits the ability of iSCAT to resolve molecular dynamics to the order of the tag size; homogeneous tags could potentially allow the resolution to be taken below this limit by deconvolution methods. In a wider context, the presented harmonic potential well confinement model has the potential to detect and characterise a wide variety of biological phenomena, such as hop diffusion, receptor clustering, and lipid rafts.

## Introduction

Single particle tracking (SPT) experiments directly observe the motion of single molecules, and hence offer a powerful method to analyse the membrane environment. For instance, detection and characterisation of heterogenous diffusion behaviours yields information on membrane structure (1, 2). However, SPT methods require the molecule of interest to be tagged with a trackable label that is imaged over a number of time steps. A number of experimental design limitations constrain the amount of information that can be extracted from such data, including spatial accuracy, temporal resolution and tracking period. New technologies are capable of extending the trajectory length whilst retaining high sampling rates and high spatial resolution. For example interferometric scattering microscopy (iSCAT) can generate very long (50,000 step) trajectories, with high spatial (*<* 2 nm) and temporal (up to 500 kHz) resolution (3–6). However, a fundamental problem that impacts on interpretation is the effect of the tag itself (7). This is particularly relevant for iSCAT microscopy as the gold nanoparticle (AuNP) tags are 20-40 nm in diameter whilst spatial resolution is estimated to be *∼*2 nm for a 20 nm AuNP (4, 5); relative movements between the AuNP and the bound GM1 will thus convolve with the movement of the GM1. For example, an iSCAT study on model membranes demonstrated both Gaussian-like and ring-like confinement events, which was ascribed to transient multivalent binding of the tag (4). Thus, in order to extend this technique to *in vivo* experiments there is a need to deconvolve the tag signature from the environment signal. Failure to achieve this separation means that interpretation of the high resolution dynamics measured by these techniques may be limited to the order of the tag’s size.

Analysis of SPT data is not straightforward primarily because of the stochastic nature of diffusion. This has led to the development of a range of statistical methods that detect deviations from Brownian motion, such as mean square displacement (MSD) (8–13), and confinement (14–19) analyses. A new breed of methods model switching in the movement dynamics between various dynamic states (20–24), often within a hidden Markov chain framework (25–30). For high resolution data the latter techniques can utilise the high level of information present in the trajectory to extract detailed motion characteristics, and potentially infer underlying biophysical mechanisms.

In this paper, we develop a harmonic potential well (HPW) confinement analysis within the context of a hidden Markov model (HMM). Specifically the particle moves between two states hidden to the observer: free diffusion with (to be determined) diffusion coefficient *D*, and confinement in a HPW (centre and strength to be determined). Working in a Bayesian framework, we developed a Markov chain Monte Carlo (MCMC) algorithm to infer model parameters and hidden states from a single trajectory. We tested the algorithm on simulated data, then applied it to previously published experimental iSCAT trajectories of GM1 lipids diffusing in model membranes (4). Specifically, a (20 nm or 40 nm) AuNP was coated in cholera toxin B subunits (CTxB) by streptavidin binding, each CTxB then binding 5 GM1 molecules in the lipid membrane to form an AuNP/CTxB/GM1 complex. In trajectories of 20 nm AuNP/CTxB/GM1 diffusing in model membranes on a glass substrate, we detected clear periods of trapping in wells of mean radius 18nm with a mean trapping time 0.024s. However, we also observed inherent heterogeneities in both AuNP/CTxB/GM1 particles and trapping sites, which ultimately affect trajectory characteristics.

This paper is organised as follows. In Methods we introduce the HPW confinement model and an associated inference (MCMC) algorithm. The full derivation of the MCMC algorithm is described in Note S1 in the Supporting Material. In Results we demonstrate accurate inference of model parameters and hidden states on simulated trajectories, then apply the algorithm to iSCAT trajectories of AuNP/CTxB/GM1 diffusing in model membranes.

## Methods

### Harmonic potential well model

We developed a model for a particle that switches between a freely diffusing state, and a confinement state localised around a slowly diffusing centre. The state is encoded by a hidden variable *z*, with *z*_*i*_ = 0 if the particle is freely diffusing at time *t*_*i*_, and *z*_*i*_ = 1 if confined, where *i* = 1*..N* denotes the timepoint (i.e. frame). The state *z*_*i*+1_ depends only on *z*_*i*_ with transition probabilities (constant frame rate)

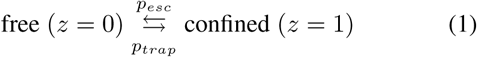

where *p*_*trap*_ and *p*_*esc*_ are the per frame probabilities of switching into and out of confinement respectively. The probability of being in state *z*_*i*+1_ given state *z*_*i*_ is therefore

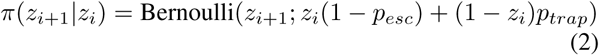

where Bernoulli(*x*; *p*) denotes the Bernoulli probability distribution with variable *x* and parameter *p*. In the free state the particle diffuses freely with diffusion coefficient *D*. In the confined state the particle is assumed to have a directed component to its diffusive motion, proportional to the distance from the well centre *C*_*i*_, i.e. the force is proportional to *X*_*i*_ *-C*_*i*_ where *X*_*i*_ is the particle position at time *t*_*i*_. (Note that *X*_*i*_ and *C*_*i*_ are 2D vectors.) During confinement the centre diffuses much slower than the particle itself (diffusion coefficient *D*_*C*_ *<< D*). When the particle is free *C* diffuses with diffusion coefficient *D*_*est*_, where *D*_*est*_ is sufficiently high that the centre can relocate between different confinement sites. The centre is thus still present even when it is not affecting the particle. The stochastic differential equations (SDEs) for this model are

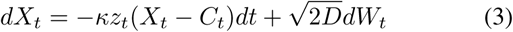

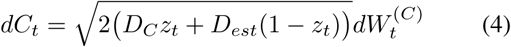

where 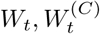 are independent Weiner processes. During confinement *X*_*t*_ has Ornstein-Uhlenbeck (OU) dynamics with centre *C*_*t*_. We assume that switching can only occur at the sampling points. We also assume that *C*_*t*_ is slowly varying and therefore ignore its time dependence over the time step Δ*t*. The frame-to-frame dynamics are hence

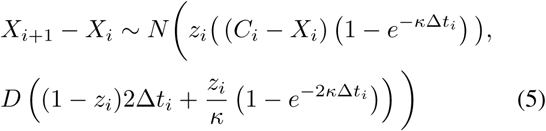

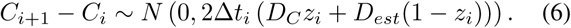

See Note S1 for full details. If the step size is sufficiently small relative to the confinement strength (*κ*Δ*t <<* 1) an Euler-Maruyama approximation is justified, but if the particle explores the well over Δ*t*, this OU solution is required. We refer to this discrete time stochastic model as the harmonic potential well (HPW) confinement model.

The model has two hidden states to be inferred at all trajectory timepoints *i* = 1,.., *N* : the state *z*_*i*_ (confined or free), and the position of the HPW centre *C*_*i*_ when confined. There are also five parameters to be inferred: two diffusion coefficients (*D* and *D*_*C*_), the strength of the HPW (*κ*), and two transition probabilities (*p*_*esc*_ and *p*_*trap*_). *D*_*est*_ is treated separately as it only weakly affects the trajectory, and doesn’t affect the likelihood or parameter estimates provided it is sufficiently high. Fig. 1A shows a simulated HPW model trajectory. In the simulation we include a drift term for the centre so that *C* tracks *X* when not confined. This ensures that *C* is close to *X* when the particle switches from free diffusion to confinement, and therefore confinement zones remain within a reasonable field of view. This tracking of *X* by *C* isn’t included in the inference algorithm as diffusion alone is sufficient to allow the Markov chain to find high probability paths.

**Figure 1:**
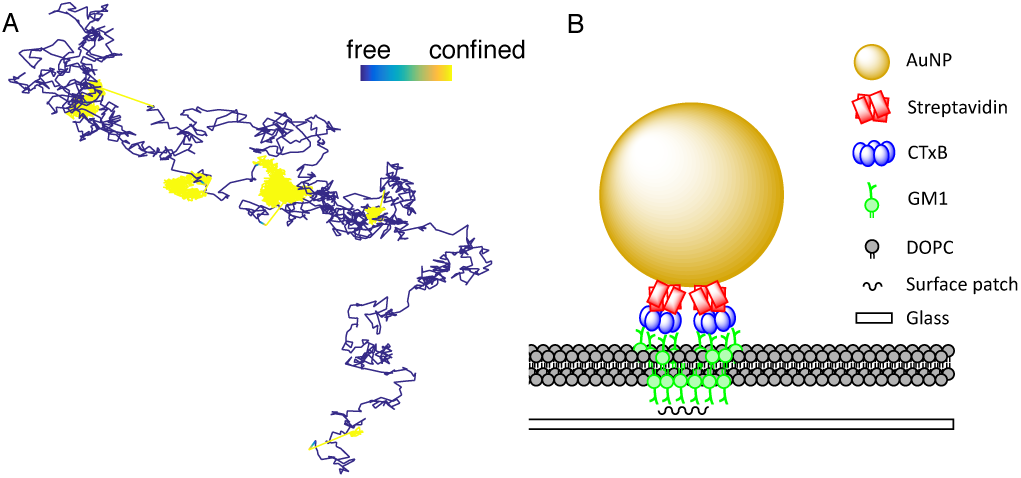
Simulated harmonic potential well (HPW) model trajectory. (A) Simulated trajectory colored by state. Model parameters: *D* = 0.5 µm^2^ s^−1^, *D*_*est*_ = 0.5 µm^2^ s^−1^, *D*_*C*_ = 0.01 µm^2^ s^−1^, *κ* = 3000 s^−1^, *p*_*esc*_ = 0.001, *p*_*trap*_ = 0.002, time step 2 × 10^4^ s, *N* = 5000 frames. Simulation performed using Equation (5), and a modified version of Equation (6) as discussed in the main text. Trajectory colour: blue free diffusion, yellow confined. Colorbar length 0.1 µm. (B) Schematic of AuNP/CTxB/GM1 complex in DOPC lipid bilayer, based on figure in reference (4).

### MCMC sampler

There are a number of MCMC samplers for linear switching models in the literature; the main distinction is whether variables are integrated out using an inverse Wishart prior (31), or a Markov chain incorporating all variables is used. The latter approach allows greater control of prior information, including use of uninformative priors, whilst the Wishart distribution, motivated by computational convenience, imposes a dependence between variable correlations and scale which is a concern for inference (32). We developed an MCMC algorithm (Note S1) for the full system of variables to fit the HPW model (Equations 5, 6) to 2D trajectory data 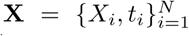. We chose unin-formative priors for all parameters except for the transition probabilities, where we use an informative prior to restrict rapid switching between states (details in Note S1). For an SPT trajectory, the algorithm samples the posterior distribution, *π*(*θ*, **z**, **C**|**X**), giving *K* samples of the parameters 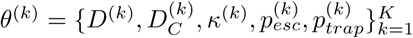, and hidden states 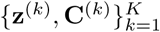. Here for each sample *k*, 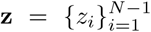 and 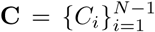 are the set of hidden states and centre locations (2D vectors) throughout the trajectory.

We determined convergence of the MCMC sampler by calculating the Gelman point scale reduction factor (PSRF) (33), considering a run converged provided the PSRF was below a threshold in all variables, set to 1.2 on experimental trajectories. The MCMC run length was increased up to a maximum of 4 × 10^5^ steps on trajectories that failed the convergence criteria on shorter runs.

### GM1 molecules diffusing in model membranes

We applied the MCMC algorithm to previously published iSCAT SPT data (4), where CTxB coated AuNPs were introduced to a DOPC lipid bilayer containing 0.03% GM1 lipids (Fig 1B). A confinement event corresponds to an interaction between an AuNP/CTxB/GM1 complex on the upper leaflet with a lower leaflet GM1 that is immobilised on a glass surface. This was previously referred to as “interleaflet coupling and molecular pinning” (4). Both Gaussian and non-Gaussian confinement events were observed; we investigated these events in greater detail using our HPW model.

The dataset includes 71 trajectories of 20 nm AuNP/CTxB/GM1 diffusing in a model membrane on a glass substrate, and 18 trajectories of 40 nm AuNP/CTxB/GM1 in a model membrane on a mica substrate. There is a dynamic error in the localisation accuracy at the 50 kHz sampling rate resulting in apparent superdiffusive behaviour, which we removed by subsampling down to 5 kHz (Note S2, Fig. S1, Fig. S2). We also removed trajectory artifacts due to multiple AuNPs in the focal area (Note S2). The MCMC algorithm did not converge on 5 20 nm AuNP/CTxB/GM1 trajectories (PSRF convergence criteria of 1.2) leaving a set of 66 trajectories for further analysis. MCMC runs on all 18 40 nm AuNP/CTxB/GM1 on mica trajectories converged.

### Thresholding hidden states for lifetime analysis

For each trajectory, at each time point *i*, we computed the probability of confinement *π*(*z*_*i*_ |**X**) from the MCMC posterior distribution samples. This probability distribution is concentrated near 0 and 1 on experimental trajectories (only 2.7% of the state probabilities were between 0.2 and 0.8, Fig. S3), indicating high confidence in confinement state estimates. To annotate the trajectory by state we define the Binary signal, 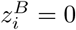 or 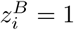, for free diffusion and confinement respectively, using a threshold of 0.5 on *π*(*z*_*i*_ **X**). We then identify confinement events as a series of ones in the (posterior) binary state vector **z**^*B*^, and free diffusions as a series of zeros, allowing event lifetimes (and per-event spatial statistics) to be computed. When considering event lifetimes we exclude those containing either the first or last timepoint of the trajectory, as the full event is not witnessed hence the state lifetime is unknown.

### Confinement event profiling

To analyse confinement events in 20 nm AuNP/CTxB/GM1 trajectories we utilised spatial statistics (including the *mean confinement radius* and *radial skewness*, defined in Table 1) based on the Euclidean distance between the particle and the confinement centre. We calculate these statistics for all events of at least 0.01 s (50 frames). Unlike the event life-time analysis, we allow events that contain either the first or last (or both) timepoints. Furthermore, we compute statistics including events revisiting a previous confinement zone where applicable (details in Table 1). These restrictions left a set of 271 confinement events when excluding repeat events, and 427 when including them. The number of events within a trajectory ranges from 1 - there were 6 examples where the particle remained trapped for the entire trajectory - to 11 (without repeats) or 25 (with repeats).

**Table 1:**
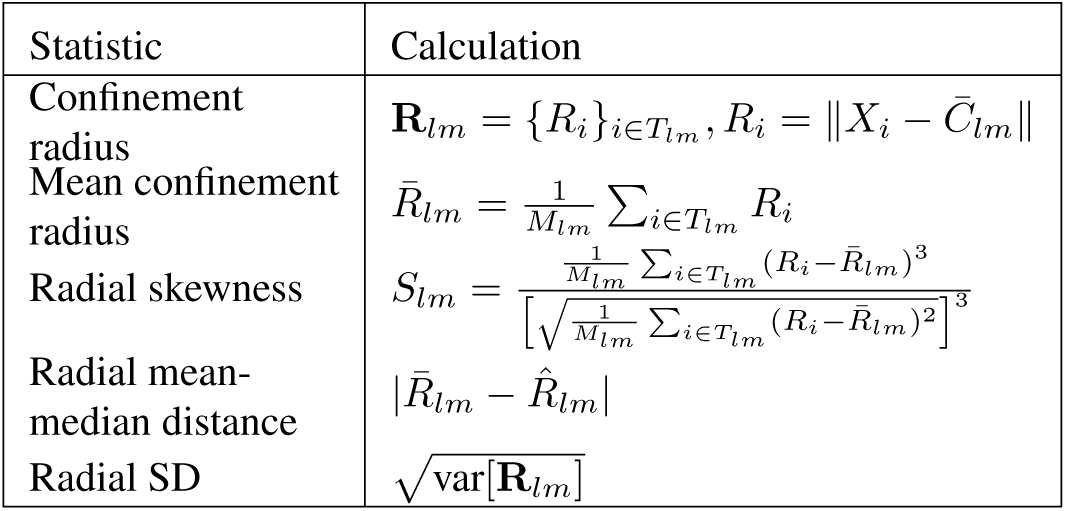
Calculation of confinement event statistics. *We denote the timepoints of the mth trapping event in the lth trajectory T_lm_. Events have associated particle positions 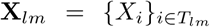 and harmonic well centre positions 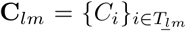. The mean posterior harmonic well cen tre is given by 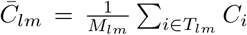, where M*_*lm*_ *is the number of timepoints in T*_*lm*_. *To remove events which revisit a previous trapping zone we didn’t include events if 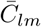was within* 30 nm *of a previous confinement centre 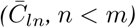 within trajectory l. ‖ . ‖ denotes the Euclidean distance, and 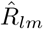 denotes the median.*

## Results

### MCMC on simulated data

The HPW model sampler was extensively tested on simulated data. Figs. 2 and 3 show an MCMC run on the simulated trajectory of Fig. 1A. The parameter posteriors are consistent with the true (i.e. simulation) values, Fig. 2. When confined, the inferred centre closely tracks the simulated centre (Fig. 3A,B), and every confinement event is accurately inferred (Fig. 3C). The inferred model parameters are independent of the algorithm parameter *D*_*est*_ (Fig. S5); although the informative priors on *p*_*trap*_, *p*_*esc*_ mean these parameters are typically underestimated. Performance was also robust to changes in trajectory length and number of events (Fig. S6, Fig. S7). Estimation of *D, D*_*C*_, *κ*, **z**, and switching rates were robust to the time series subsampling rate (Fig. S8, Fig. S9), in particular most events were still detected even with a 10 fold subsampling (Fig. S9).

**Figure 2:**
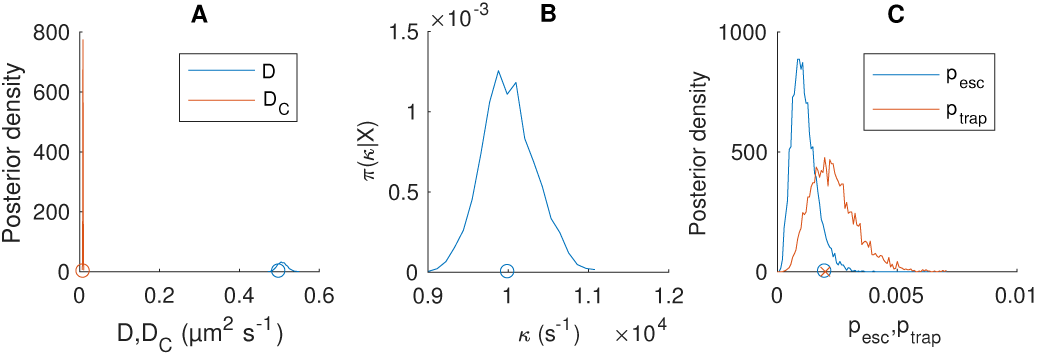
Posterior parameter distributions of the HPW model for a simulated trajectory. (A) Posterior distribution for *D* (blue) and *D*_*C*_ (red), with simulation values indicated (circles), (B) Posterior for *κ* and simulation value (circle), (C) Posterior for *p*_*esc*_ (blue) and *p*_*trap*_ (red), with simulation values (circles). Trajectory as Fig. 1. MCMC priors as Note S1. Corresponding MCMC runs shown in Fig. S4. Data based on pooling of 5 independent chains of 2000 steps with a 1000 step burn in. MCMC priors as Note S1.

**Figure 3:**
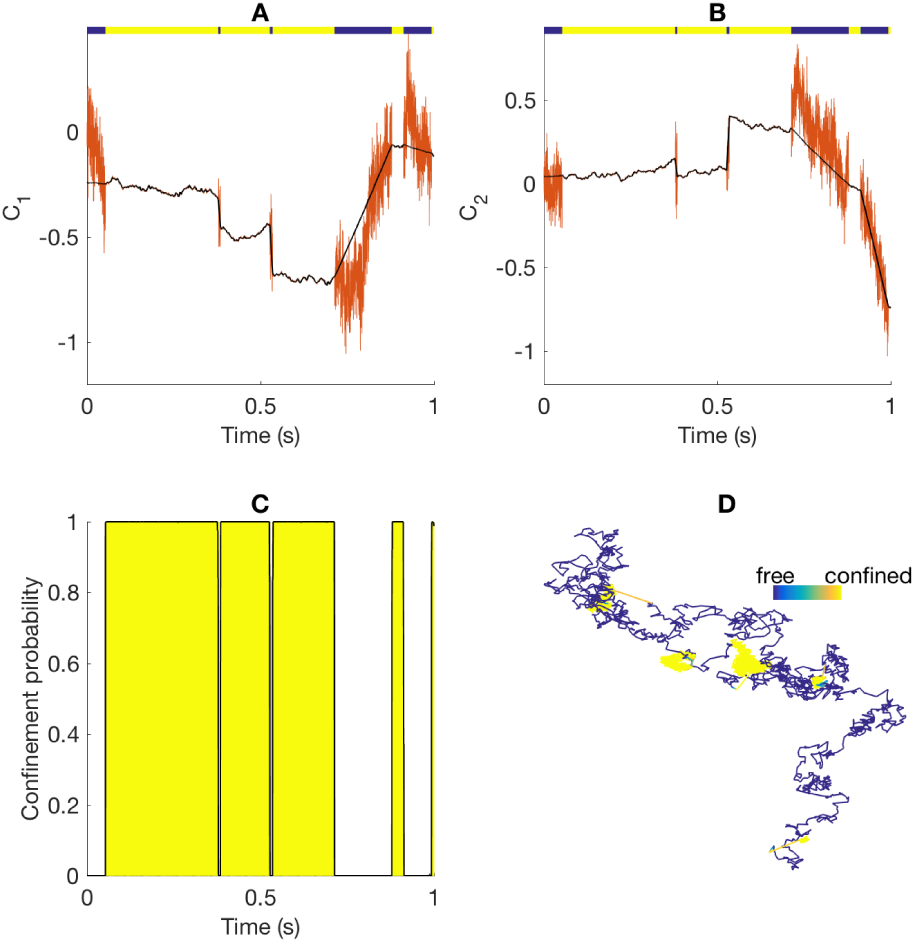
Hidden state inference for the HPW model for a simulated trajectory. (A-B) Mean inferred position of the harmonic potential centre in x and y directions (black), and simulated (true) centre (red). Coloured line at the top represents particle state: free diffusion (blue) and confinement (yellow). (C) Inferred confinement probability (black line) and simulated (true) confinement state (yellow area). (D) trajectory coloured by mean inferred confinement state, from *π*(*z*_*i*_ |**X**) = 0 (blue, free) to *π*(*z*_*i*_ |**X**) = 1 (yellow, confined). Colorbar length 0.1 µm. MCMC as Fig. 2.

### MCMC on 20 nm AuNP/CTxB/GM1 on glass trajectories

An example of the model fit is shown in Fig. 4 on the segmented trajectory shown in Fig. 4E. Fig. 5 shows the associated parameter posterior estimates. Particle state (confined or free diffusion) is well determined, with state probabilities near 0 or 1, Fig. 4C. The parameter estimates for *D* and *κ* are tight (low relative standard deviation), while the diffusion coefficient of the centre is very low, *D*_*C*_ =.010*±*0.0009 µm^2^ s^−1^ (mean*±*SD) compared to *D* =0.52 *±* 0.017 µm^2^ s^−1^, indicating near complete immobilisation of the well. The inferred position of the well centre is also practically stationary in both coordinates during periods of confinement consistent with immobilisation, Fig. 4A,B. As an independent measure of changes in mobility, we estimated the effective local diffusion coefficient (Fig. 4D), which demonstrates a clear shift at around 0.5 s (i.e. the first inferred switch point). By colour coding the trajectory according to the probability of being confined per frame, Fig. 4E, we can extract periods of confinement with non-Gaussian occupation profiles (Fig. 4F-H). In this trajectory we observed that one confinement zone is visited twice,

**Figure 4:**
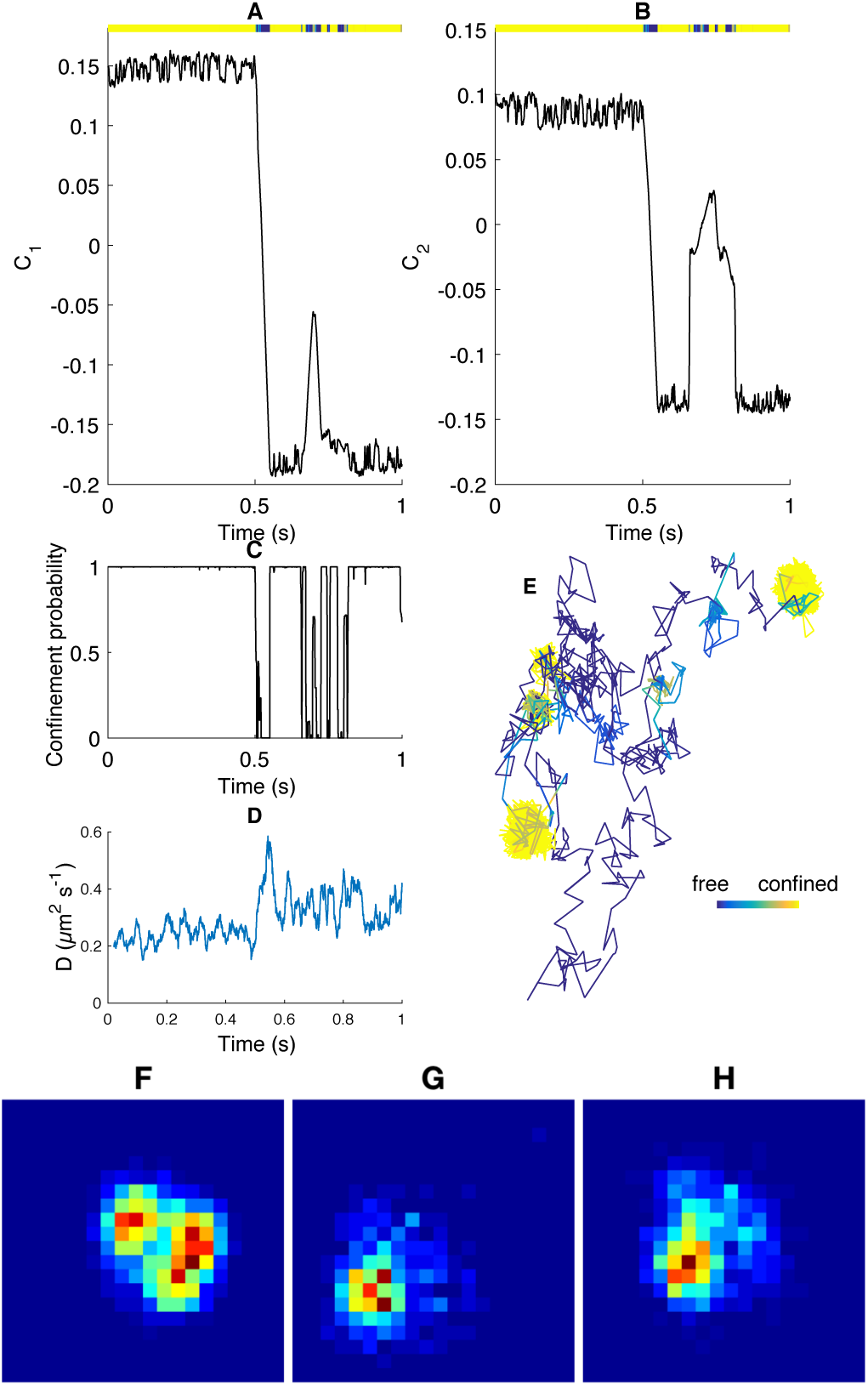
Hidden state inference for the HPW model applied to a 20 nm AuNP/CTxB/GM1 trajectory. (A-B) Mean inferred position of the harmonic potential well centre *C* (*x, y* components) and upper coloured bar representing *π*(**z** |**X**), (colour scale goes from *π*(*z*_*i*_ |**X**) = 0 (blue, free) to *π*(*z*_*i*_ |**X**) = 1 (yellow, confined)). (C) Inferred mean confinement state, (D) moving average of local maximum like-lihood diffusion coefficient estimate, window size 100 (sub-sampled frames), (E) trajectory coloured by mean inferred confinement state, colorbar length 0.1 µm. (F-H) Density coloured 2D spatial histograms of confinement events. The two events in (G) and (H) are spatially co-located. Data based on pooling of 10 independent chains. MCMC priors and convergence criteria as Note S1.

**Figure 5:**
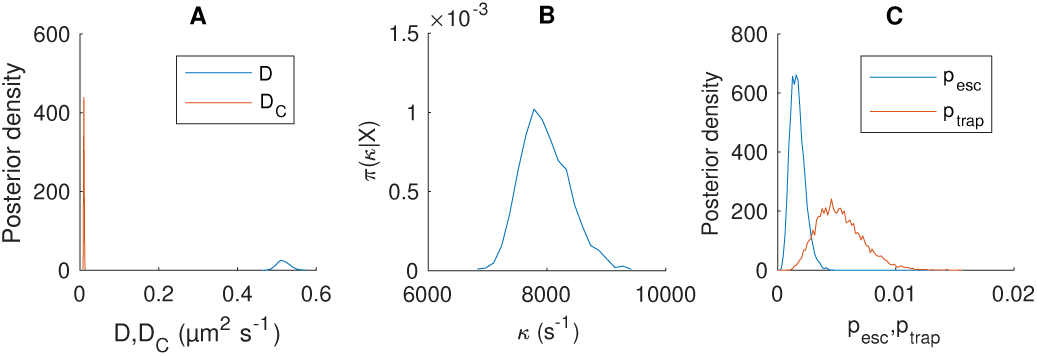
Posterior parameters of the HPW model applied to a 20 nm AuNP/CTxB/GM1 trajectory. (A) Posterior distributions for *D* (blue) and *D*_*C*_ (red). (B) Posterior for *κ*. (C) Posteriors for *p*_*esc*_ and *p*_*trap*_. Distributions consist of samples pooled from 12 independent runs. Corresponding MCMC chains shown in Fig. S10. Trajectory and MCMC runs as Fig. 4.

Fig. S11, and that the repeat confinements had remarkably similar occupation profiles, Fig. 4G-H. The probability per frame of switching is reasonably well inferred (Fig. 5C) despite the small number of events. The probability of escape from a confinement zone is smaller than the probability of trapping, reflecting the short periods of time that the AuNP/CTxB/GM1 complex is undergoing free diffusion.

Applying our MCMC algorithm to the 66 trajectories we obtain parameter estimates across the population, Fig. S12. The mean value of *D* over all trajectories was 1.15 *±* 0.106 µm^2^ s^−1^ (mean *±* SEM, population SD 0.86 µm^2^ s^−1^); MSD analysis (using the @msd-analyzer package (34)) gave a smaller estimate, 0.0525 *±* 0.017 µm^2^ s^−1^ (mean *±* SEM). This difference reflects the fact that MSD doesn’t account for confinement, which is the dominant state, whereas our diffusion coefficient estimate does. Our Bayesian analysis provides estimates of parameter confidence per trajectory which are in fact substantially smaller than the spread between trajectories, Fig. S13; specifically the ratio of the population variances of *D* and *κ* are 257 and 59 times larger than the average trajectory posterior variances respectively. This indicates the presence of system variability, giving rise to trajectory heterogeneity. To understand its cause, we investigate whether heterogeneity is manifest in the confinement events of individual trajectories, specifically the size, shape and lifetime of these events.

### Lifetime and shape analysis of confinement events

The mean confinement state lifetime (as defined in Methods) is 0.024 s, but there is a large variation in event lifetimes across trajectories (Fig. 6A-B), and significant heterogeneity across trajectories (*p* = 0.02, Kruskal-Wallis test, 1779 events across 60 trajectories). The lifetimes of free diffusion events, mean 0.002 s, did not show significant heterogeneity across trajectories, (*p* = 0.86, Kruskal-Wallis test on 60 trajectories, 1770 events). Further, we examined if the population of lifetimes across trajectories conform to an exponential waiting time model, i.e. whether switching between states obey first order kinetics. A Q-Q plot demonstrates that there is a distinct deviation from an exponential distribution fit (mean event time *µ* = 0.024s), specifically there are a far higher proportion of longer trapping events, indicative of heterogeneity. A mixture of 2 exponentials is a better fit, Fig. 6D, suggesting that the confinement events derive from a heterogeneous population with at least two components with short and long average lifetimes. The minor population of long lifetime events are dispersed over trajectories, Fig. 6B, in particular trajectories are not split into two groups with long and short mean confinement times. In contrast the free diffusion state lifetimes closely follow a mono-exponential distribution, Fig. 6E-F.

**Figure 6:**
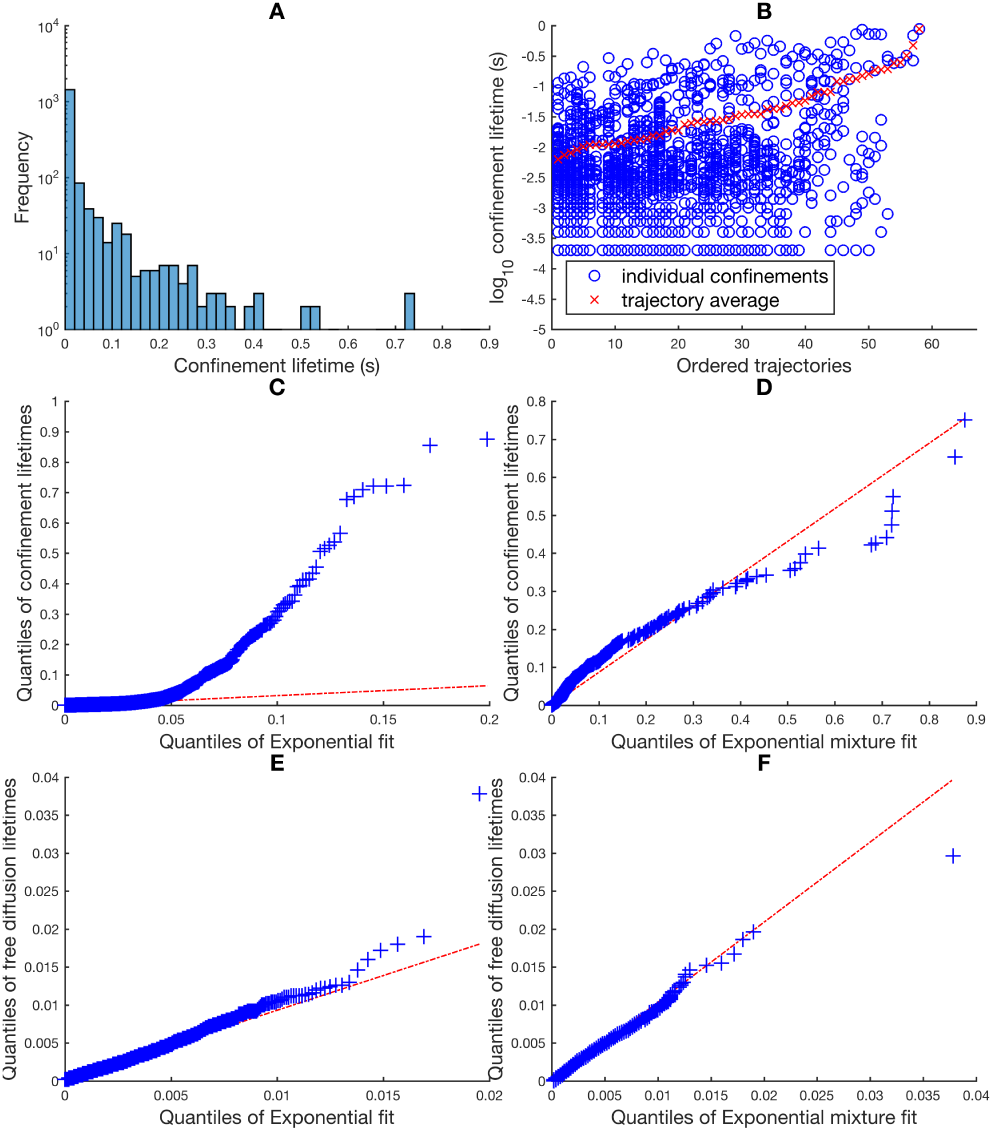
Confinement event lifetimes are not exponentially distributed. A) Histogram of all confinement lifetimes (n=1959 events). B) Scatterplot of confinement lifetimes against trajectories ordered by mean confinement lifetimes. (C-F) Q-Q plots of state lifetimes against exponential fits.C) Confinement events against the exponential distribution (*µ* = 0.024 s, *R*^2^ = 0), D) confinement events against samples (*n* = 10^4^) from mixture of 2 exponentials (*µ*_1_ = 0.004 s, *µ*_2_ = 0.1 s, weights 0.80 and 0.20 respectively, *R*^2^ = 0.98). E) Free diffusion lifetimes (2011 events) against the exponential distribution (*µ* = 0.002 s, *R*^2^ = 0.98), F) free diffusion lifetimes against samples (*n* = 10^4^) from mixture of 2 exponentials (*µ*_1_ = 0.002 s, *µ*_2_ = 0.01 s, weights and 0.01 respectively, *R*^2^ = 0.98). Red line is extrapolated linear fit to the first and third quantiles. Plots A-F include all confinement events except those which contained the trajectories’ first or last timepoint.

We next analysed confinement event shape using the spatial statistics defined in Table 1. The mean confinement radius over all trajectories is 18 nm, comparable to the size of the AuNP, although estimator inflation is likely to be present (35). The mean radial skewness is 0.88; for comparison a 2D Gaussian distribution gives a radial displacement (from the mean) that is Rayleigh distributed with skew 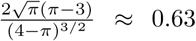. Mean confinement radius and radial skewness show a wide distribution of values across confinement events (Fig. 7A,B), with significant (one-way ANOVA: mean confinement radius *p* = 1.2 × 10^−6^, radial skewness *p* = 9.1 × 10^−4^; 271 events grouped by 66 trajectories) heterogeneity across trajectories, Fig. 7C,D. Confinement event spatial histograms for all 66 20 nm AuNP/CTxB/GM1 trajectories, ordered by the average within-trajectory mean confinement radius (Fig. S14), and average radial skewness (Fig. S15) demonstrate the wide variety of confinement shapes.

**Figure 7:**
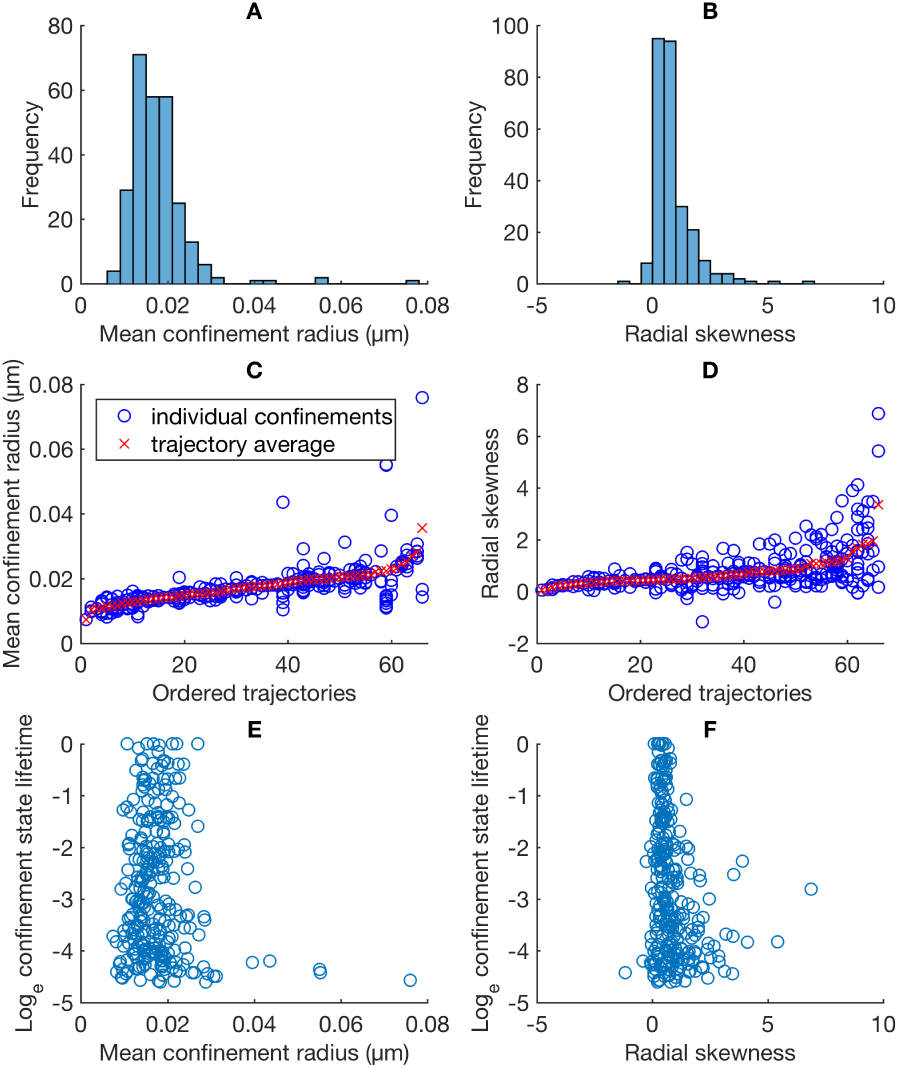
Shape statistics for confinement events in 20 nm AuNP/CTxB/GM1 trajectories. (A-B) Histograms over confinement events. (C-D) Spatial statistics for all confinement events, ordered by the average within trajectory statistic. Plots include all confinement events of at least 0.01s, with events revisiting a previous trapping zone removed (giving 271 events). (E-F) Scatterplots against trajectories ordered by average within trajectory value of the statistic.

The observed heterogeneity in confinement time, size and (previously reported (4)) shape raises two key questions:

1. Does the shape of confinement events determine their lifetime?
2. Does heterogeneity predominately arise from a mechanism operating at individual confinement sites (local environment dependent), or at the level of trajectories (AuNP/CTxB/GM1 nanoparticle dependent)?

To probe the relationship between confinement event shape and lifetime (question 1), we examined their correlation. We found no correlation between confinement state lifetime and mean confinement radius (Table S1, Fig. 7E), and a weak (but significant) negative correlation between lifetime and radial skewness, (Table S1, Fig. 7F). This suggests that the mixed-exponential nature of the binding lifetime is only weakly related to the shape of the binding event, i.e. these arise from different physical mechanisms.

Regarding question 2, the heterogeneity analysis above (see also Fig 7C,D) indicates that confinement events are statistically more similar within trajectories than across trajectories. Additionally, the ratio of the mean variance within trajectories to the variance across all events is 0.6 for both mean confinement radius and radial skewness, Table S2. To determine which confinement statistic is most strongly conserved within trajectories, we clustered events by each confinement statistic and quantified the similarity of events within single trajectories, Figure 8. Confinement size is the most conserved, followed by confinement lifetime when excluding events revisiting a previous confinement zone, Figure 8A. Incorporating revisiting events dramatically improves the conservation of confinement size and shape statistics relative to lifetime, Figure 8B; this suggests that, whilst shape is conserved, confinement time is variable between events at the same location. This shape conservation at the same site is evident from the confinement event spatial histograms for long events, Figure 9. Of note, the mean-median distance statistic failed to show significant heterogeneity across trajectories, p=0.06, one-way ANOVA, reflecting its lack of conservation when excluding events revisiting a previous confinement zone, Figure 8A, but was as conserved as life-time when all events were included, Figure 8B. Thus, in answer to question 2, heterogeneity arises at both the trajectory and confinement site level, with different effects on confinement lifetime and shape. This suggest that nanoparticle confinement events are described by 2 degrees of freedom.

**Figure 8:**
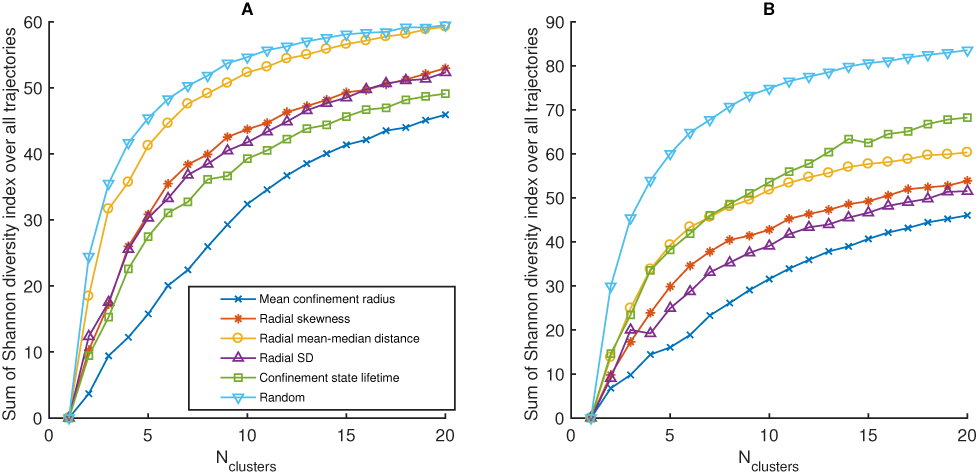
Clustering of confinement event statistics. Individual confinement events were clustered (k-means++ algorithm (36) with squared Euclidean distance metric) based on event statistics. For each trajectory, *l*, the Shannon diversity index, 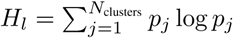 was calculated (*p*_*j*_ is the proportion of the events in trajectory *l* that appeared in cluster *j*). The sum of the Shannon diversity index over all trajectories is then a measure of the dissimilarity of events within trajectories (the lower the Shannon index the higher the similarity). The event statistics (shown in the legend) are defined in Table 1. For each choice of *N*_clusters_, 50 separate clusterings were performed (since the k-means++ algorithm stochastically assigns initial values for cluster centroids). and the sum of the Shannon diversity index was averaged over these clusterings. A) Clustering of events obtained as described in “confinement event profiling” in the main text, except with events containing the first or last timepoints excluded (214 events total). B) Same as (A), except with events which revisited a previous confinement zone included.

**Figure 9:**
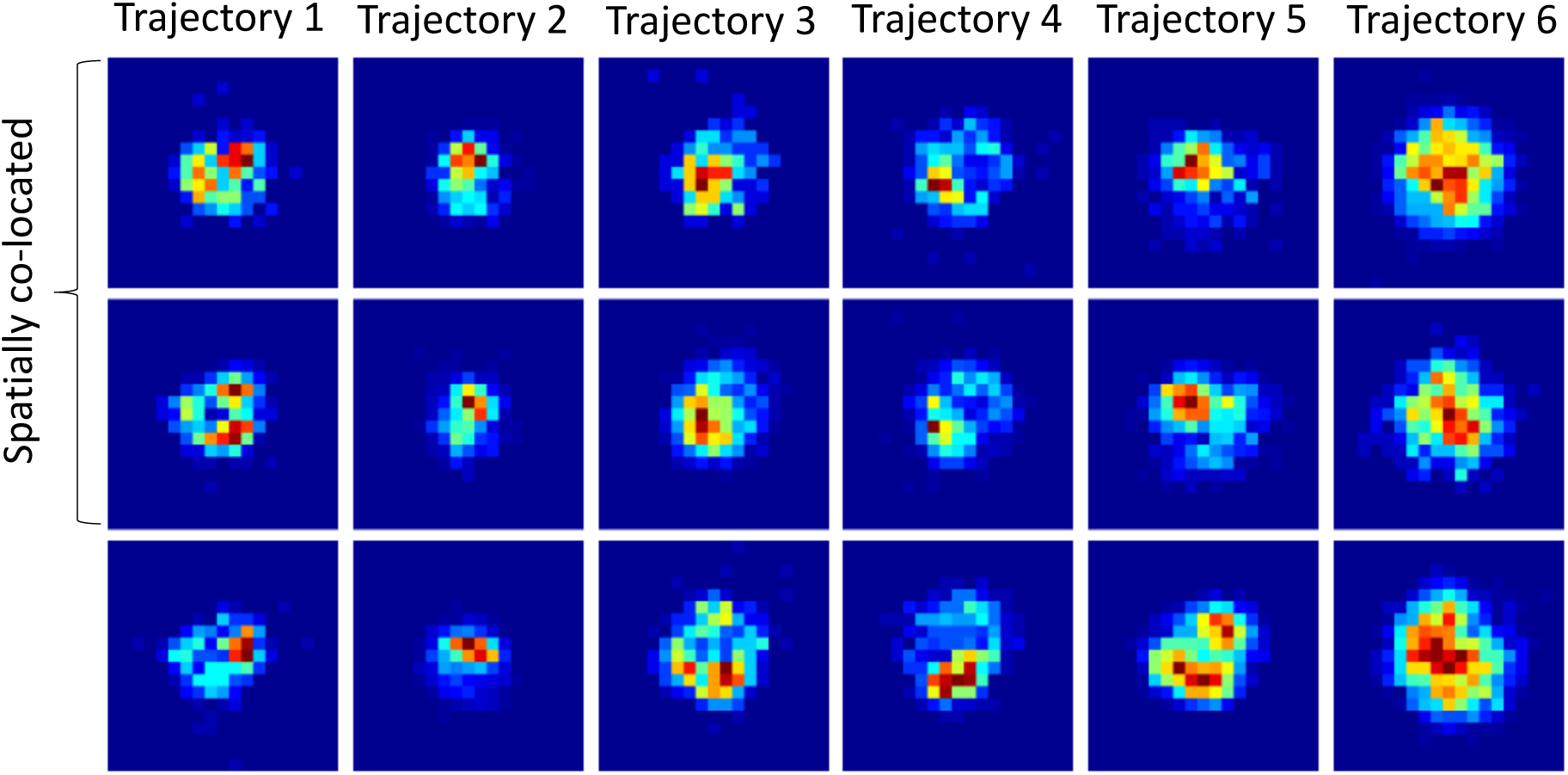
Spatial conservation of confinement events revisiting the same site. Each column shows particle position histograms for two spatially co-located confinement events, and one event at a different location in the same trajectory. The spatially co-located confinement events are distinct - i.e. the particle moved away from the trapping zone between the displayed events. Each plot has side length 0.1 µm.

### Analysis of 40 nm AuNP/CTxB/GM1 trajectories on mica

As a control, we analysed 18 trajectories of 40 nm AuNP/CTxB/GM1 diffusing in SLBs on a mica substrate. The previous analysis demonstrated that no confinement for this treatment was present (4). We applied our HPW model MCMC algorithm to this data and detected no confinement events (Fig. S16) - the posterior confinement probability was *<* 0.01 for all time, in all trajectories. The mean *D* was 1.2048 *±* 0.09 µm^2^ s^−1^ (mean *±* SEM), comparable to 20 nm AuNP/CTxB/GM1 on glass trajectories (1.15 *±* 0.20 µm^2^ s^−1^). The mean MSD-derived (with @msdana-lyzer (34)) *D* was 0.87 *±* 0.12 µm^2^ s^−1^. These values are in closer agreement than for the 20 nm AuNP/CTxB/GM1 on glass dataset, which is expected due to the lack of confinement. However, we again observed trajectory heterogeneity in the diffusion coefficients with a ratio of population variance to mean trajectory variance of 123, indicative of individual AuNP dependent diffusion coefficients.

## Discussion

We developed a Bayesian algorithm to infer a HPW confinement HMM, and used it to partition SPT trajectories into periods of free diffusion and confinement. When applied to experimental AuNP/CTxB/GM1 trajectories we detected clear periods of confinement and free diffusion, Fig. 4. It was previously proposed that confinement event shape heterogeneity (Gaussian versus non-Gaussian confinement) in this dataset was due to transient multivalent binding of the tag (4). Our analysis of confinement events attained using the HPW model revealed the following heterogeneity trends:

- Confinement size and shape are conserved within trajectories (Fig. 7C, D), and repeat events at the same site show similarities (Fig. 5G, H and Fig. 9).
- Confinement event lifetimes are heterogeneous across trajectories and comprise a mixture of at least two Exponentials with short (4 ms) and long (100 ms) mean lifetimes (Fig. 6C,D).
- Spatial heterogeneity and lifetime heterogeneity are effectively uncorrelated, suggesting they arise from different mechanisms.

Based on these observations we propose a refinement to the transient multivalent tag binding hypothesis. Namely, the characteristics of individual confinement events are determined by two factors: the size and geometry of the GM1 platform on the lower leaflet (determining residence times), and the number and distribution of CTxB complexes bound to the surface of the AuNP (determining size and shape of confinement event), Fig 1B.

These dependencies are consistent with CTxBs remaining attached to the surface via GM1s throughout the entire trajectory (Fig. 10); this is supported by the high affinity of the CTxB/GM1 bond with a dissociation rate in SLBs of (2.8 *±* 0.1) × 10^−4^ s^−1^, giving a mean binding life-time of 3.6 × 10^4^ s (37). We propose that differences in the geometry of bound CTxB on the surface of the nanoparticle causes trajectory-conserved variation in the observed confinement as follows: tightly packed (or single) CTxBs have more freedom to “wobble” (Fig. 10B), and broadly spaced, multiple (bound) CTxBs having less freedom (Fig. 10C), giving a large, respectively small confinement radius for binding events. Additionally, non-Gaussian confinement events occur when there is a second (or potentially multiple) CTxB/GM1 attachment that is not immobilised, which restricts movement to a rotation or non-uniform “wobbling” around the immobilised binding site (Fig. 10D). These hypotheses are consistent with the fact that there are around 25 CTxB per 20 nm AuNP (4) - it is expected that there will be variability in both their number and spatial distribution. We observe a strong correlation of the diffusion coefficient with the mean confinement radius (*r* = 0.61, Fig. 11 A), but not with the confinement event lifetime (*r* = 0.27, Fig. 11 B). This is consistent with the hypothesis that with more attachments the AuNP experiences higher drag, whilst the mean confinement radius decreases because of stronger geometric constraints. Our analysis thus suggests that variation in the number and spatial configuration of bound CTxB contributes to the nature of the AuNP interaction with the upper leaflet of the bilayer, thereby giving each individual AuNP/CTxB/GM1 complex a confinement signature (Fig. 8, Fig. S14 and Fig. S15), and diffusion coefficient - the latter also being evident in the confinement-free 40 nm AuNP data.

**Figure 10:**
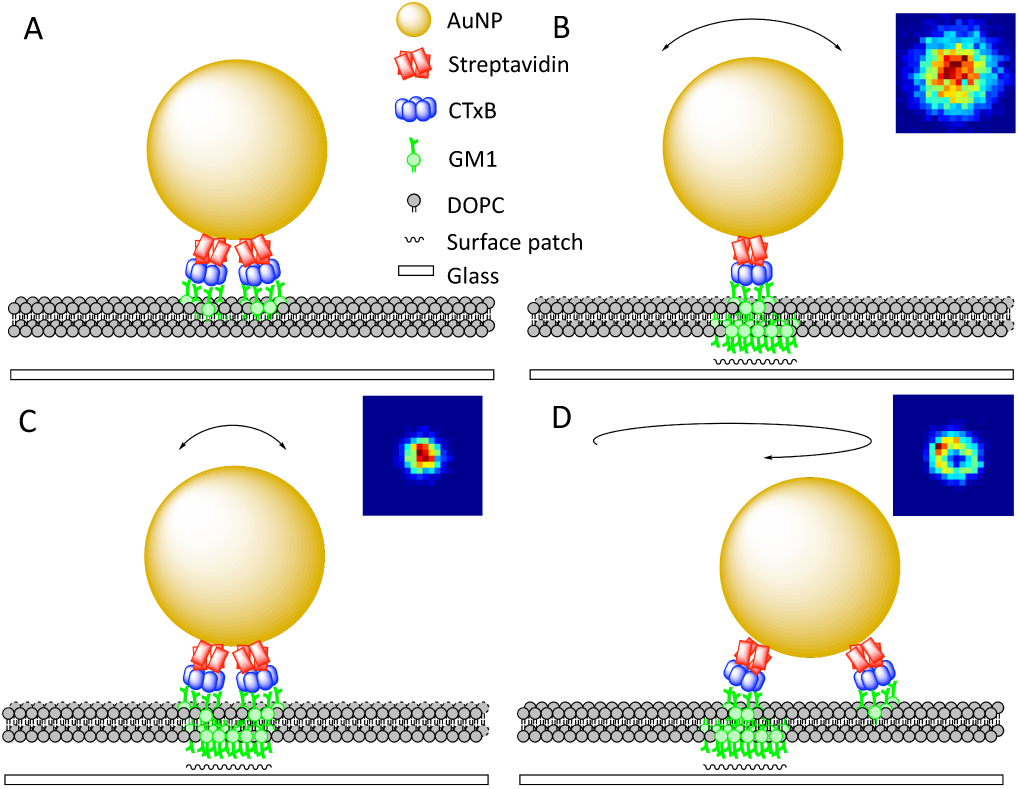
Schematic of AuNP/CTxB/GM1 structures leading to Gaussian and non-Gaussian confinement profiles. (A) Free diffusion, (B) wide Gaussian-like confinement, (C) narrow Gaussian-like confinement, (D) non-Gaussian confinement. Insets in (B-D) are example histograms of particle positions pooled over confinement events within selected trajectories (e.g. Fig. S14, Fig. S15). Insets have side length 0.1 µm. Schematic based on a figure in reference (4).

**Figure 11:**
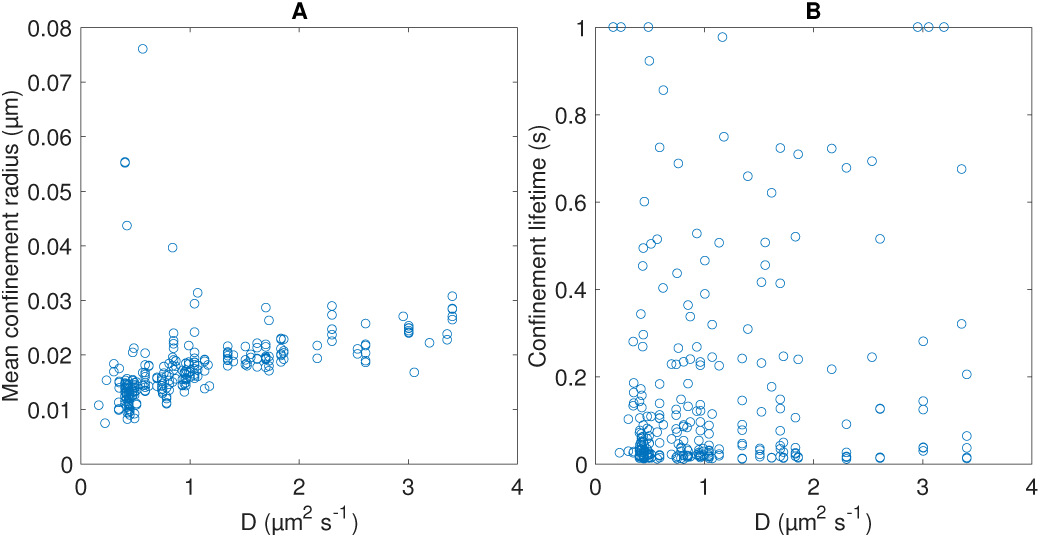
Correlation between D and confinement statistics. (A) mean confinement radius, (B) confinement lifetime. Plots include all confinement events of at least 0.01s.

The characteristics of the lower leaflet GM1 platform contribute a confinement site dependence, with conservation of shape and size upon revisiting the same site (Figs. 8, 9). On the other hand, confinement lifetime at the same site is variable. The GM1 in the lower leaflet are immobilised by hydroxyl pinning sites on the glass surface, with these sites having an estimated size of *<*10 nm (4). However, aggregation of GM1 with domain sizes of 15-60 nm in supported lipid bilayers has been observed in AFM experiments (38). Large sites consist of more aggregated GM1 in the lower leaflet. Our mean confinement radius is 18 nm, which would comprise both AuNP/GM1/CTxB nanoparticle degrees of freedom around the binding site, and displacements of the GM1 platforms between the leaflets. This suggests that either pinning sites are small, or no relative movement is possible. Larger pinning sites may trap multiple CTxB molecules on the AuNP, leading to more Gaussian behaviour as rotational degrees of freedom are lost and possibly longer (on average) trapping times. This could be the cause of the negative correlation between non-Gaussian confinement shape and event lifetime (the lower the radial skewness statistic, the longer the typical confinement time, Fig. 7F). However, the double exponential mixture distribution of confinement lifetimes cannot be explained by these mechanisms. Mean lifetime does not partition by trajectories - long event life-times being distributed throughout the trajectories, Fig. 6B - suggesting a random process is responsible. The simplest explanation is that binding of the AuNP/CTxB/GM1 at the pinning sites is heterogeneous, e.g. there could be a multi-step binding sequence with the second (long lifetime) step proceeding in only a fraction of the binding events. We note that the 6 trajectories that remain confined throughout are a third population, since even on the long lifetime distribution, observing binding events of 1s or longer is negligible (probability 4.5 × 10^−5^).

### Outlook and future work

Analysis of SPT trajectories with HMMs has advantages over other methods for detecting confinement in single trajectories. In particular, they do not rely on tuning algorithm parameters through a comparison with Brownian motion. Additional parameters (such as the confinement strength *κ*, centre *C*, and switching times as inferred here) can also be extracted, which allows for interpretation and comparison of confinement event characteristics across and within trajectories. However, appropriate HMMs are necessary for successful analysis. Specifically, models must approximate well the behaviour of different dynamic states in the data. For instance, confinement is often associated with a decrease in the effective diffusion coefficient, suggesting that models that switch diffusivities (25–30) should also be able to detect confinement in these iSCAT particle trajectories. However, we found that a two-state diffusion coefficient switching HMM (30) could not segment these trajectories (data not shown). This implies that the effective diffusion coefficient of the AuNP/CTxB/GM1 complex doesn’t change sufficiently during confinement events. The effective diffusion coefficient under confined Brownian motion is dependent on the temporal and spatial resolution of the data, suggesting that the high temporal sampling rate and positional accuracy of iSCAT data does not reduce the effective diffusion coefficient during confinement. In fact, the sampling time scale leads to displacements that are on the order of the confinement radius which required us to use OU dynamics in our MCMC algorithm. Failing to account for the effect of the confinement frame-to-frame leads to a bias in the parameter estimates (not shown).

We incorporated centre diffusion in our HPW model to relax the constraint of a circular potential. The importance of this will depend on the spatial-temporal resolution of the data and whether the confinement zone is static, or has time-dependent shape variation or drift. For instance, transient confinement in lipid rafts - which both diffuse and have an irregular shape - has been hypothesized.

The observation that individual AuNP/CTxB/GM1 complexes have a specific spatial signature means that distinguishing the effects of the tag from other factors, such as the cell membrane environment, is difficult. It follows that homogenous tags should improve characterisation of the membrane environment. Since the variability in the tag signature presumably arises from the random placing of CTxB molecules on the AuNP surface, using particles with a structured surface is predicted to reduce or potentially eliminate this problem. Virions are ideal, given their highly geometric 3D structure. Interferometric label-free tracking of virions has been demonstrated at 3 s temporal resolution (39); thus achieving the high spatial and temporal resolution of recent iSCAT microscopy (such as in the data set explored here) with viral particle tags is a distinct possibility. The resolution of the tag’s signature and the length of events will then determine the resolution of that trajectory, and thus the length scale to which SPT can discriminate different types of particle movement. Whether this can be taken below the size of the tag, reminiscent of super-resolution, remains to be ascertained.

## Conclusion

We use a HMM-based analysis to partition SPT trajectories into periods of free diffusion and confinement. Our algorithm infers the switching times between these two states, the diffusion coefficient *D*, and the characteristics of the confinement events: the HPW strength *κ*, the position of the HPW centre *C*, and the centre diffusion coefficient *D*_*C*_. We demonstrate the utility of the method on simulated and experimental data; on simulated data confinement zones were accurately detected and HPW centres accurately tracked whilst experimental trajectories were partitioned with high confidence. The model could potentially detect various biological phenomena such as lipid microdomains (or “rafts”), receptor clustering, and hop diffusion.

## Author Contributions

Analysed the data: PJS, NJB. Wrote the paper: PJS, NJB. Developed the software: PJS.

## Acknowledgements

We thank Philipp Kukura for providing the dataset for analysis, and for helpful discussions and suggestions. We also thank Christian Eggeling and Gabrielle de Wit for helpful discussions and suggestions. Paddy Slator received funding from the University of Warwick to study at the EPSRC and BBSRC funded Warwick Systems Biology Doctoral Training Centre.

